# Calcium-associated Pro367 ring puckering supports conformational selection in a thermophilic lipase

**DOI:** 10.64898/2026.10.06.756949

**Authors:** Muhammet Sait Eyerci, Ahmet Tülek, Fatma İnci Özdemir, Gözde Şükür, Hasan DeMirci

## Abstract

Thermophilic bacterial lipases must combine structural stability with the conformational flexibility required for interfacial activation, yet the local organization of their metal-binding sites remains incompletely understood. Here, we determined two X-ray crystal structures of the family I.5 lipase from *Geobacillus kaustophilus* (*Gk*_Lip_). The 2.47-Å orthorhombic model contains a Ca²⁺ ion at the conserved metal-binding site, whereas the 2.30-Å anisotropic triclinic model contains four non-equivalent protein chains in which no Ca²⁺ ion was modeled. *Gk*_Lip_ retains the canonical α/β-hydrolase fold, the Ser114-Asp318-His359 catalytic triad, the Phe17/Gln115 oxyanion-hole region, an extended lid, and a conserved Zn²⁺-binding site. In the orthorhombic structure, Gly287, Glu361, Asp366, and Pro367 contribute to the Ca²⁺ coordination environment. Pro367 adopts a Cγ-endo pucker in the Ca²⁺-containing structure; three Ca²⁺-free chains also adopt the endo pucker, whereas one adopts a Cγ-exo pucker. The presence of both Pro367 pucker states in the Ca²⁺-free structure is consistent with calcium preferentially stabilizing a pre-existing, coordination-compatible conformation. This local organization is further supported by an Arg35-centered network of direct and water-mediated contacts. Comparison with the 94%-identical open *G. thermocatenulatus* lipase reveals substantial lid rearrangement despite strong conservation of the protein core. Together, these structures highlight localized conformational variability at the calcium-binding region within an otherwise highly conserved thermophilic lipase scaffold.

## Introduction

Lipases are serine hydrolases that catalyze ester-bond hydrolysis in long-chain triacylglycerols, commonly at lipid-water interfaces. Under water-limited conditions, the same catalytic machinery can mediate esterification, transesterification, and interesterification reactions, contributing to the broad use of lipases in food processing, detergents, pharmaceutical synthesis, and biodiesel production (1–3). Structural studies established that catalysis depends on a Ser-Asp/Glu-His charge-relay system and an oxyanion hole that stabilizes the tetrahedral intermediate, despite substantial diversity in lipase sequence and architecture (4–8).

Most characterized lipases adopt an α/β-hydrolase fold in which the nucleophilic serine lies in a sharply curved loop. In many true lipases, an amphipathic lid shields the active site in the closed state. Contact with a hydrophobic interface can reorganize the lid, expose the catalytic center, and generate a substrate-accessible hydrophobic surface (9–11). The lid consequently contributes to interfacial recognition, substrate access, chain-length preference, and catalytic performance (9, 12). Thermoalkalophilic lipases from *Bacillus* and *Geobacillus* are commonly assigned to bacterial lipase family I.5, whose members are relatively large (approximately 40–45 kDa) and contain an extended lid and characteristic metal-binding elements (13, 14). Recent work illustrates the functional breadth of bacterial and thermophilic lipases, including biodiesel-related enzyme discovery, polyester hydrolysis, solvent-tolerant and detergent-compatible biocatalysis, low-temperature engineering, and conformational control of lid activation (15–20).

Structures of the closely related family I.5 lipases from *G. stearothermophilus* strains P1 and L1 revealed a conserved α/β-hydrolase core, a long lid helix that covers the active site in the closed conformation, and distinct Ca²⁺ - and Zn²⁺-binding sites (14, 21). The Zn²⁺ site is coordinated by two histidine and two aspartate residues and is integrated into the enlarged family I.5 scaffold; perturbation of this region compromises structural integrity and thermal stability in homologous enzymes (22). The open structure of the closely related *G. thermocatenulatus* lipase (*Gt*_Lip_) showed that activation involves coordinated reorganization of approximately 70 residues rather than movement of a single rigid helix (13). Thus, family I.5 lipases combine a highly conserved catalytic and metal-binding core with substantial lid plasticity (13, 23, 24).

Ca²⁺ contributes differently to the organization of bacterial lipases. Calcium depletion can reorganize local structural elements, whereas calcium addition stabilizes several family I.5 enzymes and can enhance activity (24–29). Previous characterization of *Gk*_Lip_ showed an approximately 30% increase in hydrolytic activity in the presence of Ca²⁺ (28). In the homologous T6 lipase, reinforcement of the Ca²⁺ - and Zn²⁺-binding environments reduced local atomic mobility and improved stability (29). Proline is a plausible contributor to such local organization because its five-membered ring restricts backbone conformational space while sampling Cγ-endo and Cγ-exo puckers (30–35). Ring puckering can influence backbone geometry, folding stability, and ligand affinity (31, 32, 36, 37), but its relationship to metal occupancy in family I.5 lipases has not been structurally resolved.

*G. kaustophilus* is a thermophilic bacterium with multiple genomic adaptations to growth at elevated temperature (38). Its family I.5 lipase, *Gk*_Lip_, is an approximately 43-kDa enzyme with reported optimal activity at pH 8.0 and 50 °C (28). The enzyme retains activity under several chemically demanding conditions, and modification of its N-terminal polyhistidine tag alters substrate preference and hydrolytic activity (28). *Gk*_Lip_ shares approximately 94% sequence identity with structurally characterized *Gt*_Lip_ (13), but its three-dimensional structure and the conformational organization of its metal-binding sites have not been described.

In this study, we determined *Gk*_Lip_ structures in two crystal forms, yielding five non-equivalent molecular models. One crystal form contains a modeled Ca²⁺ ion, whereas the four molecules in the second crystal form lack modeled Ca²⁺. Comparison of these models defines local Pro367 ring-puckering heterogeneity within an otherwise conserved Ca²⁺-binding region. Together with the Arg35-centered interaction network and comparisons with open, ligand-bound homologues, these structures suggest that calcium binding preferentially stabilizes a pre-existing local conformation of the metal-binding region.

## Results

### Sequence conservation and crystallographic characterization of Gk_Lip_

*Gk*_Lip_ crystallized in two forms, hereafter designated *Gk*_Lip_P212121_ and *Gk*_Lip_P1_; data-collection and refinement statistics are summarized in Table S1. The orthorhombic *Gk*_Lip_P212121_ crystal has unit-cell parameters a = 72.86 Å, b = 74.25 Å, and c = 102.64 Å and yielded a structure at 2.47 Å resolution. The model was refined to *R_work_* and *R_free_* values of 0.18 and 0.23, respectively, and contains one *Gk*_Lip_ molecule, one modeled Ca²⁺ ion, one modeled Zn²⁺ ion, and 231 water molecules. Ramachandran analysis placed 97.0% of residues in favored regions and 3.0% in allowed regions, with no outliers.

*Gk*_Lip_P1_ crystallized in the triclinic space group P1, with unit-cell parameters a = 71.50 Å, b = 74.28 Å, c = 102.02 Å, α = 89.97°, β = 89.98°, and γ = 90.20°. Diffraction was strongly anisotropic, extending to a maximum resolution of 2.30 Å, with spherical and ellipsoidal completeness values of 37.6% and 75.9%, respectively. The asymmetric unit contains four non-equivalent *Gk*_Lip_ molecules (chains A–D). The model was refined to *R_work_* and *R_free_* values of 0.21 and 0.27, respectively, and contains four modeled Zn²⁺ ions and 46 water molecules; no Ca²⁺ ion was modeled in any chain. Ramachandran analysis placed 97.3% of residues in favored regions and 2.7% in allowed regions, with no outliers.

*Gk*_Lip_P212121_ structure was determined by molecular replacement using *G. stearothermophilus* T6 lipase (PDB entry 4X6U) as the search model. The *Gk*_Lip_P1_ structure was subsequently determined by molecular replacement using an *AlphaFold*-predicted model of *Gk*_Lip_ as the search model. Together, the two crystal forms provide five non-equivalent views of *Gk*_Lip_ for comparison of the calcium-binding region.

Sequence comparison with structurally characterized bacterial lipases showed the closest relationship to *Gt*_Lip_ (PDB entry 2W22) (13). *Gk*_Lip_ and *Gt*_Lip_ share 367 identical residues across the 389-residue sequence (94%), with 381 positions having positive substitution scores (97%) and no alignment gaps (Fig. 1). Sequence identity is lower with *Pseudomonas aeruginosa* lipase (*Pa*_Lip_; PDB entry 1EX9; 30.65%) (39) and *P. cepacia* lipase (*Pc*_Lip_; PDB entry 5LIP; 23.21%) (40). The *Gk*_Lip_ catalytic triad (Ser114, Asp318, and His359), Zn²⁺ ligands (Asp62, His82, His88, and Asp239), and residues comprising the calcium-binding region are conserved in *Gt*_Lip_ (Fig. 1). Strictly conserved residues cluster within the shared structural core and around functional regions (Fig. S1).

**Figure 1.**
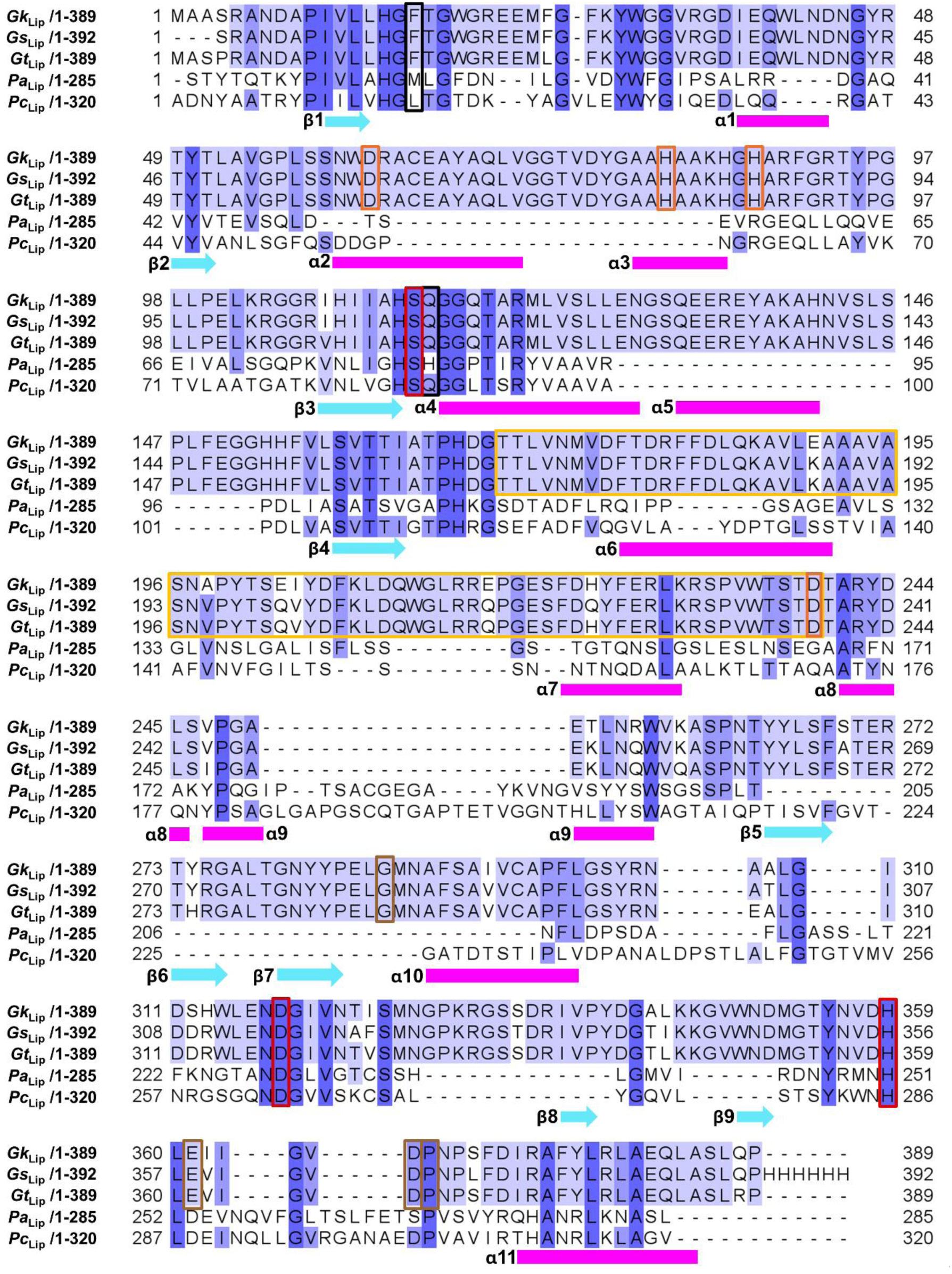
Sequence alignment of *Gk*Lip and structurally characterized bacterial lipases. *Gk*Lip was aligned with *Pa*Lip (PDB entry 1EX9; 30.65% identity), *Pc*Lip (PDB entry 5LIP; 23.21%), *Gt*Lip (PDB entry 2W22; 94.34%), and *Gs*Lip (PDB entry 4X6U; 94.82%). Blue shading indicates sequence conservation. Secondary-structure elements assigned from the *Gk*Lip_P212121 model are shown beneath the alignment; α-helices are magenta and β-strands are cyan. Frames identify the Zn²⁺ ligands (orange), catalytic triad (red), lid domain (dark yellow), calcium-binding region (brown), and oxyanion-hole residues (black).

### Overall structure of Gk_Lip_

*Gk*_Lip_ adopts the α/β-hydrolase architecture characteristic of family I.5 lipases (Fig. S2). The catalytic region lies within the hydrolase scaffold and is covered in part by a prominent lid comprising residues 169–239. The calcium-and zinc-binding sites occupy spatially distinct positions outside the immediate catalytic center (Fig. 2, *A* and *B*). *Gk*_Lip_P212121_ has a mean all-atom B factor of 37.0 Å² (Fig. 2, *C* and *D*). Electrostatic surface calculations show a pronounced negatively charged region adjacent to the calcium-binding site and a more mixed environment around the zinc-binding region (Fig. 2, *E* and *F*).

**Figure 2.**
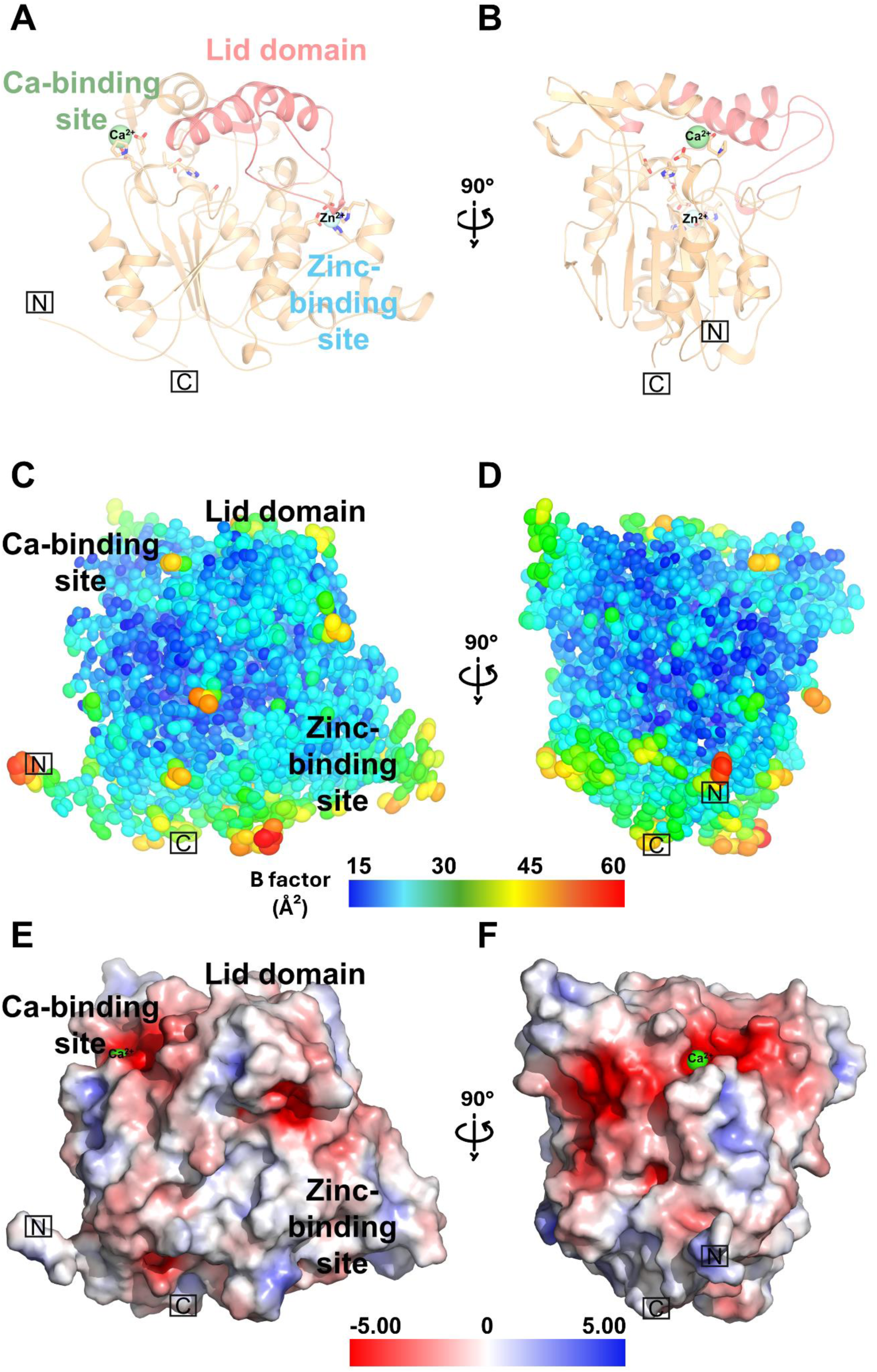
Overall structure, B-factor distribution, and electrostatic surface of *Gk*Lip. *A* and *B*, *Gk*Lip_P212121 shown in orientations related by 90°. The lid and the modeled Ca²⁺-and Zn²⁺-binding sites are indicated; Ca²⁺ and Zn²⁺ are pale green and pale cyan, respectively. *C* and *D*, B-factor representation on a 15–60 Å² blue-to-red scale. *E* and *F*, electrostatic surface potential in the corresponding orientations, colored according to the scale in the figure.

### Structural organization of the catalytic and metal-binding sites

The catalytic and metal-binding sites occupy distinct regions of the α/β-hydrolase scaffold (Fig. 3 and Fig. S3). The catalytic center comprises Ser114, Asp318, and His359, and the oxyanion-hole region is formed by Phe17 and Gln115. In *Gk*_Lip_P212121_, a Ca²⁺ ion is modeled at the conserved site formed by Gly287, Glu361, Asp366, and Pro367, with residue-to-ion distances of 2.3, 2.6, 2.8, and 2.4 Å, respectively (Fig. 3*B*). The Zn²⁺ site is formed by Asp62, His82, His88, and Asp239 (Fig. 3*D*). Continuous *2Fₒ − F*꜀ density supports the placement of the surrounding residues in the refined models (Fig. S3).

**Figure 3.**
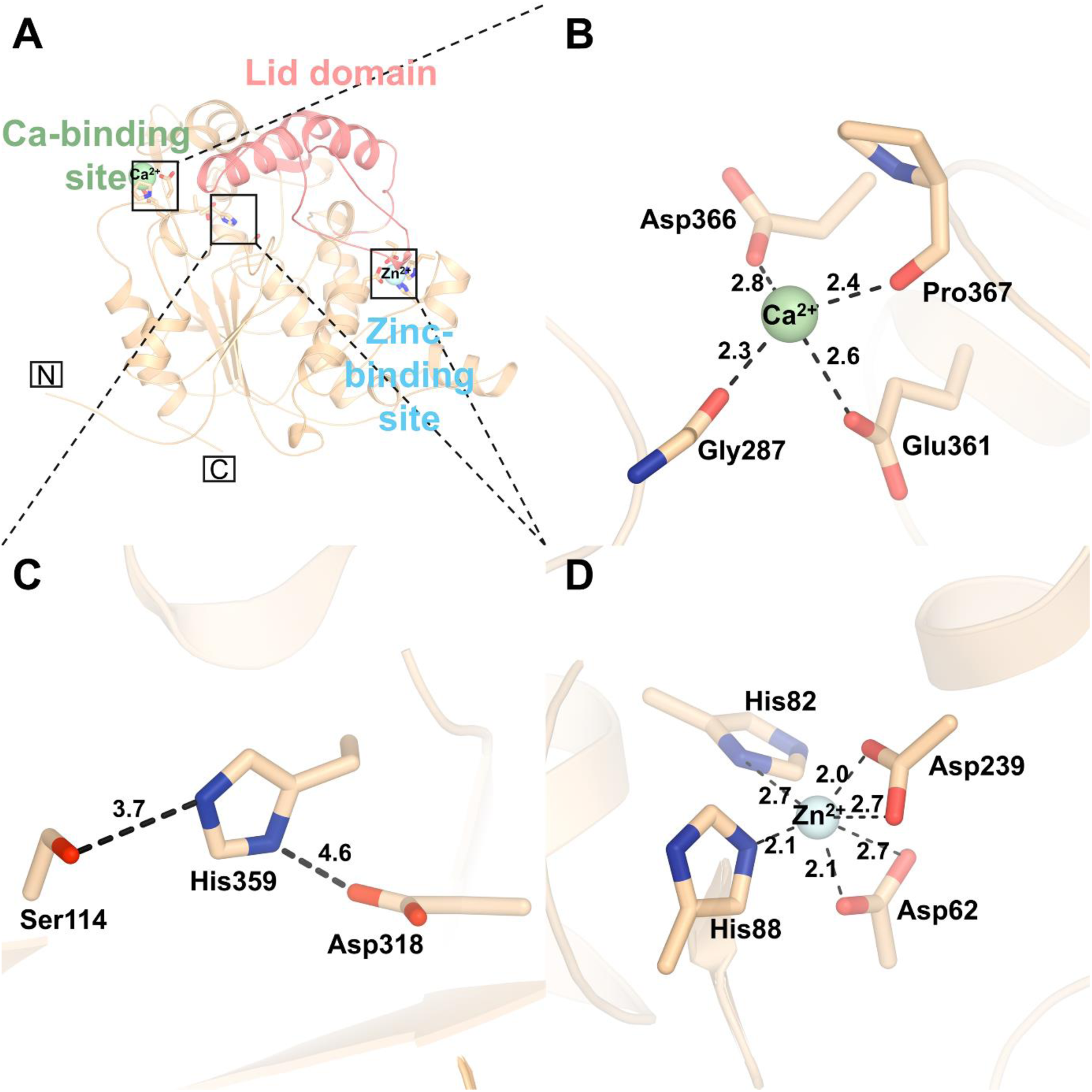
Structural organization of the principal functional sites of *Gk*Lip. *A*, *Gk*Lip_P212121 with the calcium-binding site, catalytic center, zinc-binding site, and lid indicated. *B*, calcium-binding site formed by Gly287, Glu361, Asp366, and Pro367. *C*, catalytic triad comprising Ser114, Asp318, and His359. *D*, zinc-binding site formed by Asp62, His82, His88, and Asp239. Dashed lines show the measured atom-to-ion or catalytic-site distances in Å.

### The Ca^2+^-binding region displays structural heterogeneity between Gk_Lip_ crystal forms

*Gk*_Lip_P212121_ contains a modeled Ca²⁺ ion at the site formed by Gly287, Glu361, Asp366, and Pro367, whereas no Ca²⁺ ion was modeled at the equivalent site in any *Gk*_Lip_P1_ chain. Core-based superposition shows that the five models are closely similar overall but vary locally near Pro367 (Fig. 4*A* and Fig. S4). Symmetry-aware heavy-atom RMSD values for the catalytic, oxyanion-hole, zinc-binding, and calcium-binding residues are summarized in Table S2. Within the calcium-binding region, Pro367 has the largest mean displacement across the four *Gk*_Lip_P1_ chains. The local models are supported by their respective *2Fₒ − F◻*maps (Fig. 4, *B–F*).

**Figure 4.**
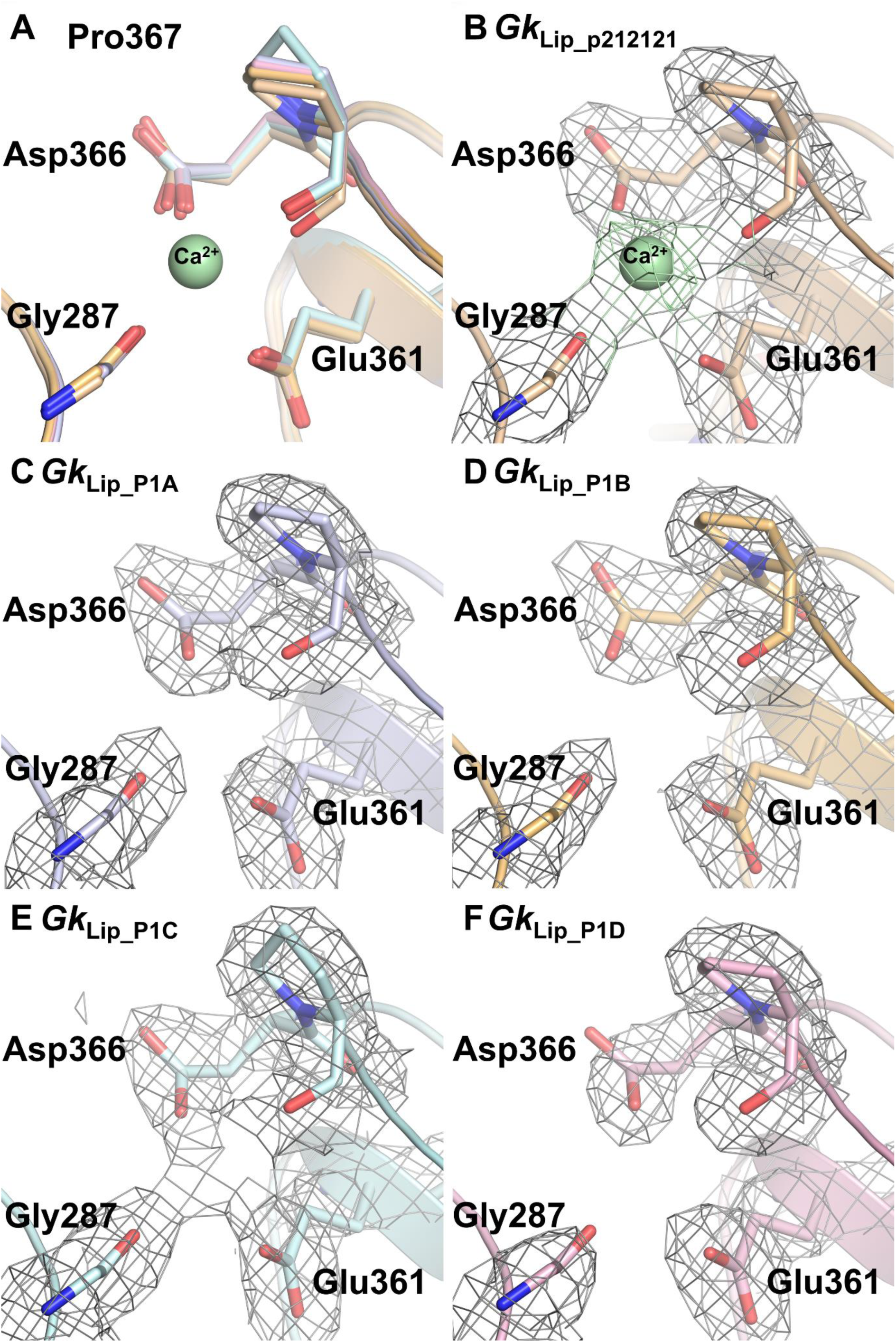
Calcium-binding-site comparison across the two *Gk*Lip crystal forms. *Gk*Lip_P212121 contains one protein chain and a modeled Ca²⁺ ion, whereas *Gk*Lip_P1 contains four non-equivalent chains without modeled Ca²⁺. *A*, superposition of the five calcium-binding regions, highlighting local variation at Pro367. *B*, *Gk*Lip_P212121, calcium-binding region. *C–F*, corresponding regions of *Gk*Lip_P1 chains A–D. Glu361, Asp366, and Pro367 are shown with their *2Fₒ − F*꜀ maps contoured at 1σ.

### Pro367 puckering heterogeneity and an Arg35-centered interaction network characterize the Ca^2+^-binding region

Backbone comparison of residues 3–389 gave mean RMSDs of 0.275–0.282 Å between the four *Gk*_Lip_P1_ chains and the *Gk*_Lip_P212121_ reference, with larger deviations confined to discrete regions (Fig. S4). In the calcium-binding site, the mean symmetry-aware heavy-atom RMSDs were 0.228 ± 0.062 Å for Gly287, 0.176 ± 0.071 Å for Glu361, 0.303 ± 0.025 Å for Asp366, and 0.516 ± 0.129 Å for Pro367 (Table S2), identifying Pro367 as the most variable of the four site residues.

Pyrrolidine-ring analysis revealed two Pro367 puckers. Pro367 is Cγ-endo in *Gk*_Lip_P212121_, with a pseudorotation phase angle of 40.27° and a puckering amplitude of 34.66°. *Gk*_Lip_P1_ chains A, B, and D are also Cγ-endo, with phase angles of 33.96°, 34.50°, and 36.71° and amplitudes of 33.45°, 33.34°, and 32.61°, respectively. By contrast, *Gk*_Lip_P1_ chain C is Cγ-exo, with a phase angle of 238.12° and an amplitude of 34.79° (Table S3). Thus, both pucker states occur in the Ca²⁺-free crystal form, whereas the Ca²⁺-containing structure adopts the endo state.

After independent core-based superposition, the Pro367 Cγ atom lies 4.9, 4.6, and 4.8 Å from the transferred Ca²⁺ position in the endo *Gk*_Lip_P1_ chains A, B, and D, respectively, compared with 4.3 Å in *Gk*_Lip_P212121_. In the exo *Gk*_Lip_P1_ chain C, the corresponding distance is 5.7 Å, approximately 0.9 Å greater than the mean for the three Ca²⁺-free endo chains (Table S3). Because no Ca²⁺ ion is present in the *Gk*_Lip_P1_ models, these values are geometric reference distances rather than coordination distances.

The Pro367 backbone carbonyl contributes to the modeled calcium-coordination environment, placing the pyrrolidine ring adjacent to the coordinating backbone geometry. The surrounding site contains an Arg35-centered network of direct and water-mediated contacts (Fig. 5). Water W30 bridges Arg35 to the Glu361 and Asp366 backbone carbonyls, with distances of 2.8 Å to Arg35 NH1, 2.9 Å to Glu361 O, and 2.7 Å to Asp366 O. Water W5 lies 3.3 Å from Arg35 NE and 3.1 Å from the Asn368 backbone N. Arg35 additionally contacts the Asn368 and Phe371 backbone carbonyls. These interactions link Arg35 to the Glu361–Phe371 segment surrounding the calcium site.

**Figure 5.**
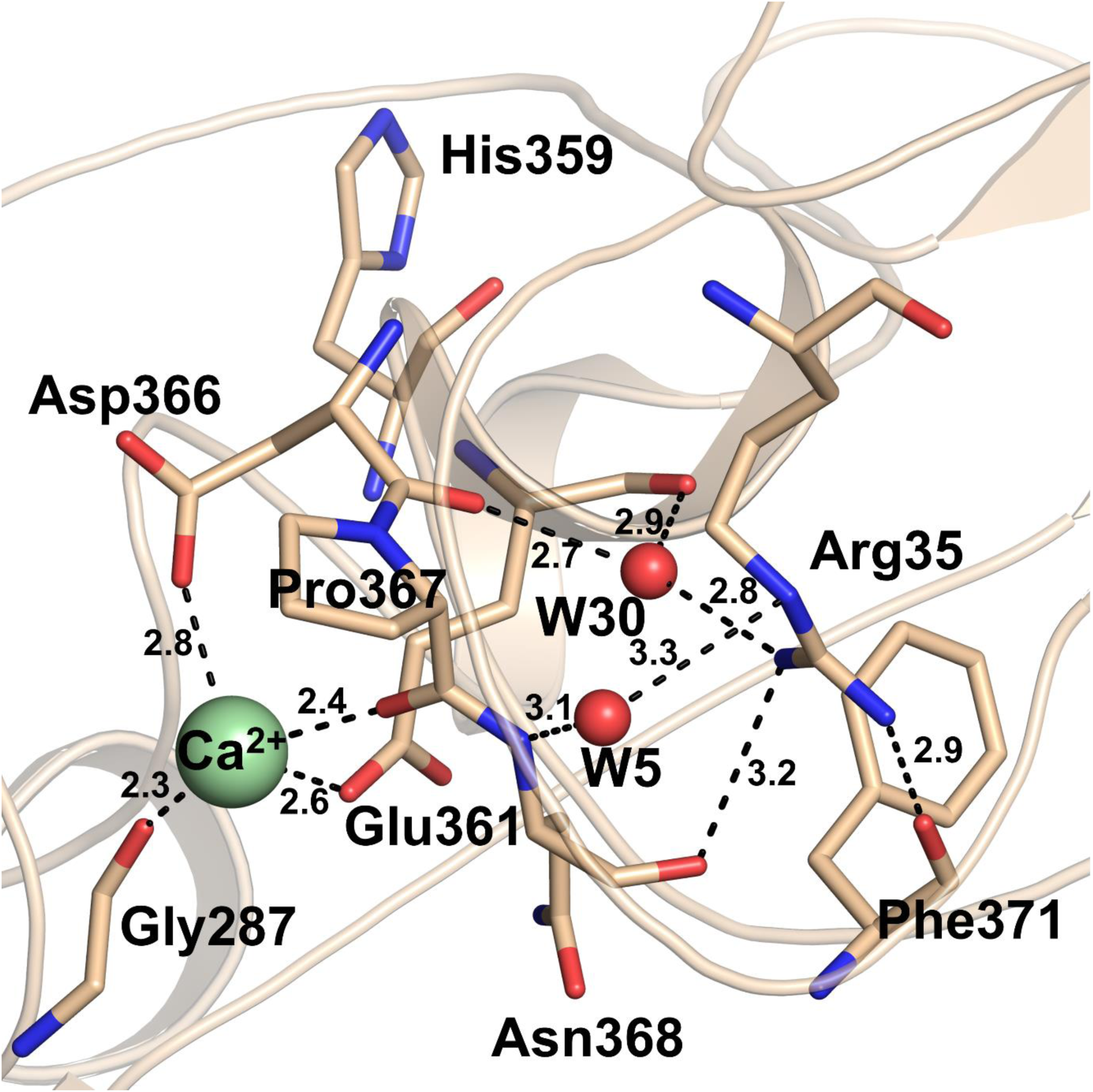
Arg35-centered interaction network at the calcium-binding site. Close-up of *Gk*Lip_P212121. The modeled Ca²⁺ ion is pale green, and waters W30 and W5 are red. W30 connects Arg35 to the Glu361 and Asp366 backbone carbonyls; W5 participates in contacts involving Arg35 and Asn368. Arg35 also contacts the Asn368 and Phe371 backbone carbonyls. Dashed lines denote measured contacts in Å. Catalytic His359 is shown to indicate its proximity to the network.

## Discussion

### Calcium-associated Pro367 puckering supports conformational selection

The occurrence of both Cγ-endo and Cγ-exo Pro367 conformers in Ca²⁺-free *Gk*_Lip_P1_ indicates that the unoccupied site can sample more than one local geometry. Notably, three of the four Ca²⁺-free chains adopt an endo pucker that falls within the same pseudorotation basin as the Ca²⁺-containing structure. This pattern is consistent with calcium binding favoring or stabilizing a pre-existing endo conformation that is compatible with the metal-binding site. The exo conformer observed in *Gk*_Lip_P1_ chain C further highlights the local flexibility of this region in the absence of calcium. Because the structures derive from different crystal forms, crystal packing and local ion availability may also influence the conformations observed.

This coupling can be considered in terms of the local geometry of the Ca²⁺-binding region. Pro367 shows the greatest variability among the four site residues, while its backbone carbonyl contributes directly to the calcium coordination environment. In *Gk*_Lip_P212121_, the Glu361–Pro367 segment is also integrated into an Arg35-centered network of direct and water-mediated contacts (Fig. 5). Calcium binding may therefore help stabilize this segment in a more defined local arrangement, constraining the geometry of the surrounding backbone, side chains, and ordered solvent.

This interpretation is consistent with studies of homologous lipases in which Arg35, ordered water molecules, and the corresponding Ca²⁺-binding loop contribute to structural stability and resistance to solvent inactivation (29). A related role for calcium in organizing loops adjacent to the active site has also been described in *P. aeruginosa* lipase (39). In *Gk*_Lip_, catalytic His359 lies immediately upstream of the Glu361–Pro367 segment and remains closely conserved among the structures.

Stabilization of this neighboring region could therefore help maintain the local environment surrounding His359 without requiring a substantial shift of the catalytic residue. This interpretation is also consistent with the previously reported increase in *Gk*_Lip_ activity in the presence of calcium (28).

### Comparison with Gt_Lip_ supports alternative lid conformations within a highly conserved scaffold

The 94%-identical *Gt*_Lip_ provides a close structural reference for the *Gk*_Lip_ lid (Fig. 1). Whole-protein superposition gives a Cα RMSD of 4.43 Å across 388 residue pairs, but this value is dominated by the lid. Excluding residues 169–239, the remaining 317 Cα atoms align with an RMSD of 0.72 Å. After this core-based superposition, the 71-residue lid regions have a Cα RMSD of 11.50 Å. The proteins therefore retain nearly identical cores while differing markedly in lid position, consistent with comparisons among other family I.5 lipases (23). *Gk*_Lip_ adopts a closed configuration in which the lid covers the catalytic region, whereas ligand-bound *Gt*_Lip_ represents an open configuration (13, 24) (Fig. 6). These structures are homologous endpoints and should not be interpreted as direct snapshots of a *Gk*_Lip_ opening trajectory. Nevertheless, localization of the major displacement to the lid supports its expected role in regulating access to the conserved catalytic triad and oxyanion-hole region.

**Figure 6.**
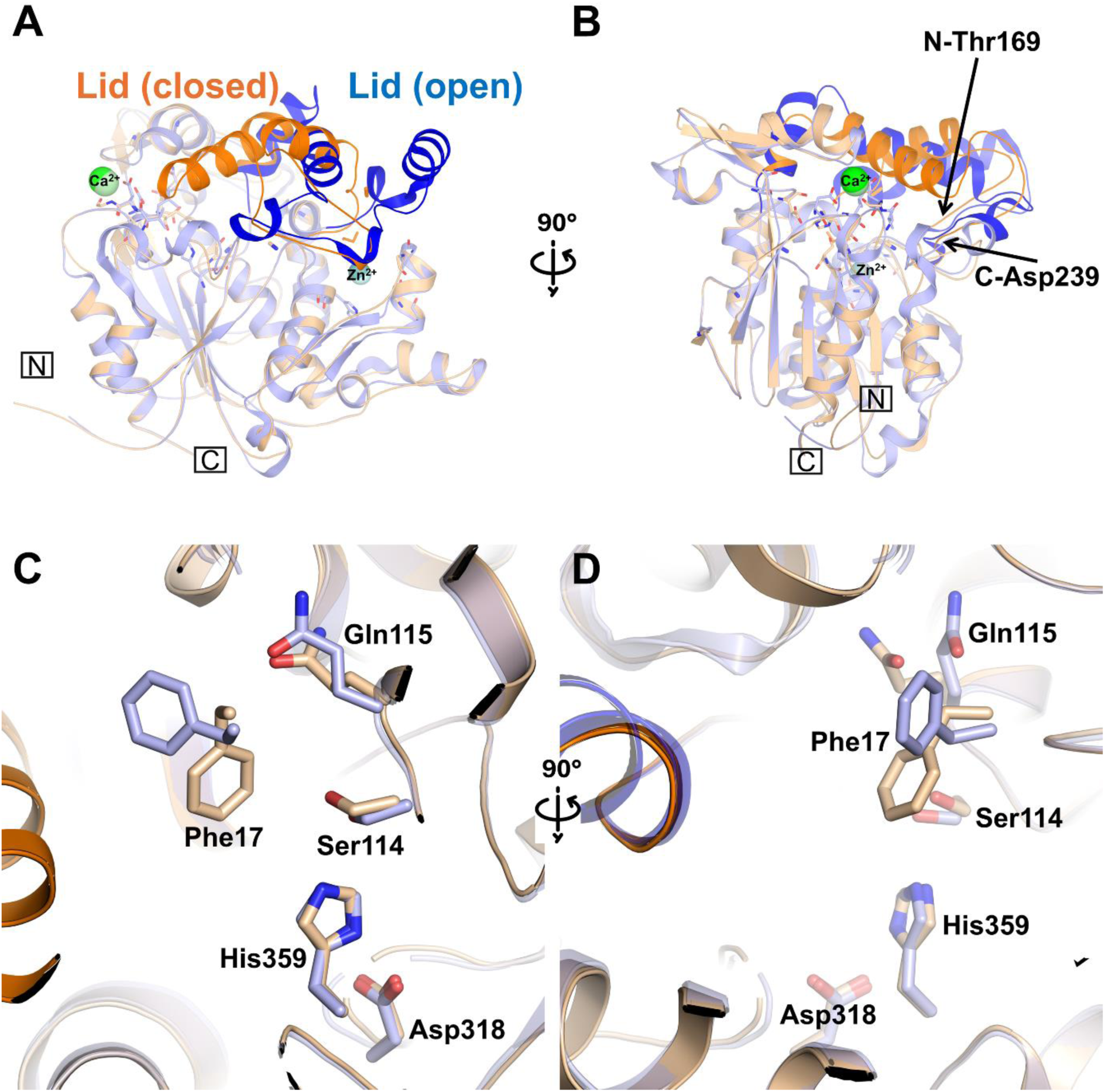
*Gk*Lip and *Gt*Lip retain a conserved core but differ in lid conformation. *Gk*Lip_P212121 was compared with open, ligand-bound *Gt*Lip (PDB entry 2W22; 94.34% sequence identity). *A* and *B*, superposition in two orientations. *Gk*Lip is wheat with an orange lid; *Gt*Lip is light blue with a blue lid. Core superposition excluding residues 169–239 gives a Cα RMSD of 0.72 Å across 317 residue pairs; after this fit, the lids differ by 11.50 Å across 71 residue pairs. *C* and *D*, catalytic region showing Phe17, Gln115, Ser114, Asp318, and His359.

### Ligand-bound homologues provide structural references for potential substrate accommodation in Gk_Lip_

No substrate-like ligand was observed in either *Gk*_Lip_ structure. To outline possible ligand-accessible space, experimentally observed ligands from homologous lipases were transferred to the *Gk*_Lip_ frame by structural superposition (Fig. 7). The closest reference, *Gt*_Lip_ (PDB entry 2W22), contains two Triton X-100 molecules in its open active-site region (13). Additional comparisons used the OCP transition-state analog in *Pa*_Lip_ (PDB entry 1EX9) and *Pc*_Lip_ (PDB entry 5LIP) (39, 40). Studies of family I.5 architecture additionally show that residues surrounding the structural zinc site can influence local packing and enzymatic behavior (41). The *Bh*_Lip_ comparison used tetradecyl hydrogen (R)-(3-azidopropyl)phosphonate (PDB ligand 1QY; PDB entry 4KE8) (42).

**Figure 7.**
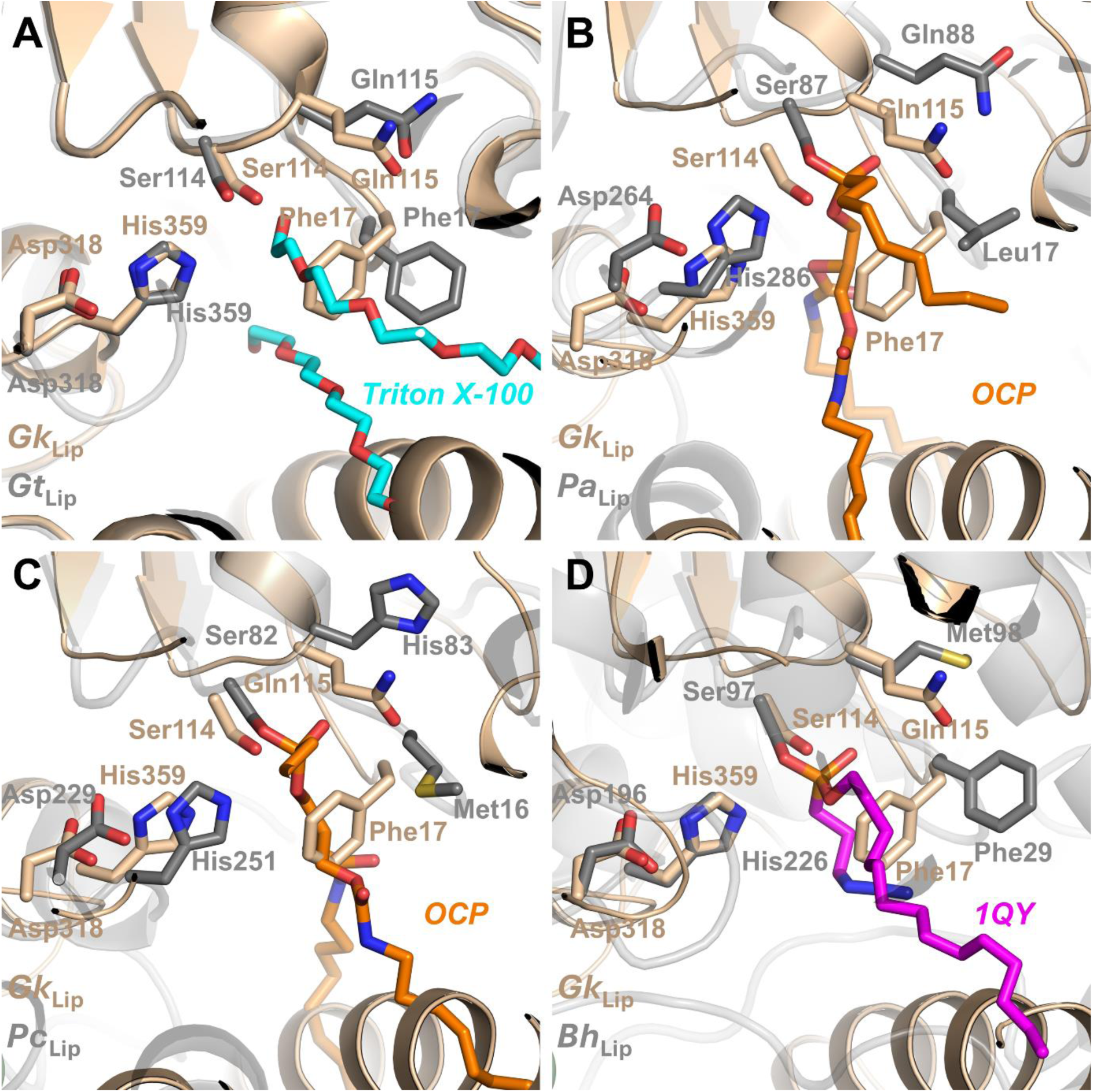
Ligand-accessible space projected from ligand-bound lipase homologues. *Gk*Lip_P212121 (wheat) was superposed with ligand-bound lipases, and the experimentally observed ligands were displayed in the *Gk*Lip frame. The *Gk*Lip catalytic triad and oxyanion-hole residues are shown. *A*, Triton X-100 from *Gt*Lip (PDB entry 2W22), cyan. *B*, OCP transition-state analog from *Pa*Lip (PDB entry 1EX9), orange. *C*, OCP from *Pc*Lip (PDB entry 5LIP), orange. *D*, tetradecyl hydrogen (R)-(3-azidopropyl)phosphonate (1QY) from *Bh*Lip (PDB entry 4KE8), magenta. These transferred poses are structural projections rather than experimentally determined *Gk*Lip binding modes.

The transferred ligands converge near the *Gk*_Lip_ catalytic triad and oxyanion-hole region but diverge as their hydrophobic groups extend toward the lid and surrounding channels. This pattern is consistent with conservation of proximal catalytic geometry and variation in distal substrate trajectories (9, 12, 43). Related crystallographic and simulation studies likewise emphasize family-dependent lid plasticity and interface-coupled conformational change (44, 45). Because the ligands were observed in other proteins and were not refined against *Gk*_Lip_ diffraction data, the transferred poses are structural projections, not experimentally determined *Gk*_Lip_ binding modes. The *Gt*_Lip_ comparison is the most informative because of the exceptionally high sequence and core-structure conservation.

## Conclusion

The two crystal forms of *Gk*_Lip_ provide five non-equivalent structural views of a highly conserved family I.5 lipase scaffold and reveal localized conformational heterogeneity within its Ca²⁺-binding region. Pro367 adopts both Cγ-endo and Cγ-exo conformations in the Ca²⁺-free structure, whereas the Ca²⁺-containing structure adopts the endo conformation, supporting a model in which calcium preferentially stabilizes a pre-existing coordination-compatible geometry. The surrounding Arg35-centered interaction network provides an additional structural context for organization of this region. Comparison with the closely related open *Gt*_Lip_ further shows that substantial lid rearrangement can occur while the catalytic core remains largely conserved. Together, these observations provide a structural framework for understanding how local calcium-site organization and lid conformational variability are accommodated within the conserved architecture of thermophilic family I.5 lipases.

## Experimental procedures

### Protein expression and purification

*Gk*_Lip_ was produced using previously optimized procedures with minor modifications (28, 46, 47). *Escherichia coli* BL21(DE3) cells carrying pET28a(+)-*Gk*_Lip_ were grown overnight at 37 °C in LB medium containing 50 µg/ml kanamycin. Fresh 100-ml cultures were inoculated at 1% (v/v) and grown at 37 °C to an OD₆ ₀ ₀ of 0.6. Expression was induced with 0.25 mM isopropyl β-D-1-thiogalactopyranoside (IPTG) for 6 h. Cells were collected by centrifugation at 8,000 × g for 15 min at 4 °C and resuspended in 50 mM Tris-HCl, pH 7.5, 200 mM NaCl, 20 mM imidazole, 0.1% (v/v) Triton X-100, and 5% (v/v) glycerol. Cells were disrupted by sonication, and the lysate was clarified at 12,000 × g for 20 min at 4 °C. The supernatant was loaded at 2.5 ml/min onto Ni-NTA agarose equilibrated in 20 mM Tris-HCl, pH 7.5, 200 mM NaCl, and 20 mM imidazole. After washing in the same buffer, *Gk*_Lip_ was eluted with 250 mM imidazole and further purified by size-exclusion chromatography on a Superdex 300 column. The N-terminal His-tag remained intact. *Gk*_Lip_-containing fractions were pooled, concentrated with a 10-kDa molecular-mass-cutoff device, adjusted to 0.21 mg/ml, and stored at −80 °C.

### Crystallization

Initial screening used microbatch-under-oil crystallization in 72-well Terasaki plates. Purified *Gk*_Lip_ (0.21 mg/ml) was mixed 1:1 with commercial sparse-matrix conditions, and each drop was covered with 16.6 µl of paraffin oil. Approximately 3,500 conditions were screened at 20 °C. Crystals corresponding to both forms were obtained from PEG RX-II condition 40: 15% (v/v) 2-propanol, 0.1 M sodium citrate tribasic dihydrate, pH 5.0, and 10% (w/v) polyethylene glycol 10,000. Crystals were flash-cooled in liquid nitrogen for data collection.

### Data collection, processing, and structure determination

Diffraction data were collected at EMBL beamline P14 at PETRA III (DESY, Hamburg, Germany) using a wavelength of 0.976254 Å and processed with *autoPROC* (48). *Gk*_Lip_P212121_ crystals belonged to space group P2₁2₁2₁, with unit-cell parameters *a* = 72.86 Å, *b* = 74.25 Å, and *c* = 102.64 Å, and diffracted to 2.47 Å. *Gk*_Lip_P1_ crystals belonged to space group P1, with unit-cell parameters *a* = 71.50 Å, *b* = 74.28 Å, *c* = 102.02 Å, α = 89.97°, β = 89.98°, and γ = 90.20°. The strongly anisotropic *Gk*_Lip_P1_ data were treated with *STARANISO*, extending to 2.30 Å with 75.90% ellipsoidal completeness (49). Data-collection and processing statistics are given in Table S1.

The *Gk*_Lip_P212121_ structure was determined by molecular replacement in *CCP4i2* using PDB entry 4X6U as the search model (50). Manual model inspection and rebuilding were carried out in *COOT*, followed by iterative refinement with *REFMAC5* within *CCP4i2* and *PHENIX* (51−53). PHENIX refinement included individual coordinate and isotropic atomic displacement parameter refinement, TLS refinement using five groups, and occupancy refinement, with automatic optimization of coordinate and ADP weights. The final model contains the 389-residue *Gk*_Lip_ sequence, one additional His-tag-derived histidine, one modeled Ca²⁺ ion, one modeled Zn²⁺ ion, and 231 ordered water molecules. The N-terminal His tag was retained during purification and crystallization; however, only one tag-derived histidine was sufficiently ordered to be modeled in the *Gk*_Lip_P212121_ structure. The structure was refined at 2.47 Å to *R_work_* and *R_free_* values of 0.18 and 0.23, respectively. Ramachandran analysis placed 97.0% of residues in favored regions and 3.0% in allowed regions, with no outliers.

The *Gk*_Lip_P1_ structure contains four *Gk*_Lip_ molecules (chains A–D) in the asymmetric unit and was determined by molecular replacement using an *AlphaFold*-predicted model of *Gk*_Lip_ as the search model (54). Model inspection and rebuilding were performed in *COOT*, and refinement was carried out in *PHENIX* using coordinate refinement, individual isotropic atomic displacement parameters, and torsion-angle non-crystallographic symmetry restraints among the four chains. Residues 359–368, encompassing catalytic His359 and the Ca²⁺-binding segment analyzed here, were excluded from the NCS restraints to allow independent refinement of their local conformations. The final model contains four 389-residue *Gk*_Lip_ chains, four modeled Zn²⁺ ions, and 46 ordered water molecules; no Ca²⁺ ion was modeled. The structure was refined at 2.30 Å resolution to *R_work_* and *R_free_* values of 0.21 and 0.27, respectively. Ramachandran analysis placed 97.3% of residues in favored regions and 2.7% in allowed regions, with no outliers.

The Zn²⁺ site was supported by anomalous difference density and its coordination by Asp62, His82, His88, and Asp239. The ion at the conserved calcium-binding site of *Gk*_Lip_P212121_ was modeled as Ca²⁺ based on the local electron density, oxygen-donor coordination environment and distances, and correspondence with the conserved Ca²⁺ site of homologous family I.5 lipases. No anomalous signal was used to establish the Ca²⁺ assignment.

### Sequence and structural analyses

Protein sequence alignments were generated using *Clustal Omega* and visualized and annotated in *JALVIEW* (55, 56). Structural analyses, superpositions, distance measurements, and molecular graphics were performed in *PyMOL* (57).

*Gk*_Lip_ and *Gt*_Lip_ structures were first aligned across the complete protein without iterative outlier rejection. The lid was defined as residues 169–239. Core alignment excluded the lid, and the lid RMSD was then calculated without additional fitting. For comparisons between crystal forms, each *Gk*_Lip_P1_ chain was independently superposed on *Gk*_Lip_P212121_ using Cα atoms from residues 1–168 and 240–389. Residue-wise backbone RMSDs were calculated from the N, Cα, C, and O atoms without further local fitting (Fig. S4). Heavy-atom RMSDs for functional-site residues were calculated in the same structural frame (Table S2); for residues containing symmetry-equivalent atoms, the lower chemically equivalent assignment was retained. Pro367 geometry was evaluated from its five endocyclic torsion angles to obtain the pseudorotation phase angle and puckering amplitude and to classify Cγ-endo and Cγ-exo conformers (Table S3). Distances from Pro367 Cγ in the Ca²⁺-free chains were measured to the Ca²⁺ position transferred from the superposed *Gk*_Lip_P212121_ structure and therefore represent geometric reference distances rather than coordination distances.

## Data availability

All data supporting the findings are contained in the article and its Supporting Information. Coordinates and structure factors have been deposited in the Protein Data Bank under accession codes 45HU (*Gk*_Lip_P212121_) and 45IY (*Gk*_Lip_P1_). The *PyMOL* scripts used for structural comparisons, RMSD calculations, Pro367 pseudorotation analysis, and related geometric measurements have been deposited in *Zenodo* under DOI 10.5281/zenodo.22662843.

## CRediT Author Statement

Sait Eyerci: Investigation, Formal analysis, Writing - Original Draft. Ahmet Tülek: Formal analysis, Writing - Original Draft, Supervision. F. İnci Özdemir: Investigation. Gözde Şükür: Investigation. Hasan DeMirci: Formal analysis, Writing - Original Draft, Supervision.

## Supplementary material

This article contains Supplementary Material.

## Funding and additional information

This work was supported by the Scientific and Technological Research Council of Türkiye (TÜBİTAK) 2209-A program (project 1919B012506377 to M.S.E.). The funder had no role in the study design; collection, analysis, or interpretation of data; preparation of the manuscript; or the decision to submit the article for publication.

## Declaration of generative AI and AI-assisted technologies

During the preparation of this work, the authors used ChatGPT (OpenAI) to assist with language editing and manuscript organization. After using this tool, the authors reviewed and edited the content as needed and take full responsibility for the content of the article.

## Supporting information

Supplementary information including S1-S4 and Tables 1-3

## Acknowledgments

Diffraction data were collected at EMBL beamline P14 at PETRA III (DESY, Hamburg, Germany). We thank David von Stetten and Arwen Pearson for assistance with data collection.

## Abbreviations

*Bh*_Lip_: *Bacillus* sp. H257 lipase
*Gk*Lip: *Geobacillus kaustophilus* DSM 7263T lipase
*Gs*_Lip_: *G. stearothermophilus* T6 lipase
*Gt*_Lip_: *G. thermocatenulatus* lipase
Ni-NTA: nickel-nitrilotriacetic acid
OCP: phosphonate transition-state analog
*Pa*Lip: *Pseudomonas aeruginosa* lipase
*Pc*Lip: *Pseudomonas cepacia* lipase
PDB: Protein Data Bank
RMSD: root-mean-square deviation.

