## Supplementary information including S1-S4 and Tables 1-3 for "Calcium-associated Pro367 ring puckering supports conformational selection in a thermophilic lipase"

**Supporting information for "Calcium-associated Pro367 ring puckering supports conformational selection in a thermophilic lipase"**

M. Sait Eyerci [(1)], Ahmet Tülek [(2)], F. İnci Özdemir [(3)] , Gözde Şükür [(3)], Hasan DeMirci [(4)]

(1) From the Department of Molecular Biology and Genetics, Faculty of Science, Boğaziçi University, Istanbul 34342, Türkiye

(2) From the Kütahya Health Sciences University, Faculty of Engineering and Natural Sciences, Department of Molecular Biology and Genetics, Merkez, Kütahya 43100, Türkiye

(3) From the Gebze Technical University, Faculty of Science, Department of Molecular Biology and Genetics, Gebze, Kocaeli 41400, Türkiye

(4) From the Department of Molecular Biology and Genetics, Faculty of Science, Koç University, Istanbul 34450, Türkiye

**Figure S1. Three-dimensional distribution of sequence-conserved residues among *Gk*$_{Lip}$ and homologous bacterial lipases.** Structural superposition of *Gk*$_{Lip\_P212121}$ with the homologous lipases from *G. thermocatenulatus* (PDB ID: 2W22), *G. stearothermophilus* (PDB ID: 4X6U), *P. aeruginosa* (PDB ID: 1EX9), and *P. cepacia* (PDB ID: 5LIP). Protein structures are shown as semi-transparent gray cartoons following superposition onto *Gk*$_{Lip}$. Residues that are strictly conserved (identical) across all five sequences in the multiple-sequence alignment are highlighted in blue. The positions of the *Gk*$_{Lip}$ lid domain and the $Ca^{2+}$ and $Zn^{2+}$-binding regions are indicated for spatial reference. N- and C-termini are labeled.

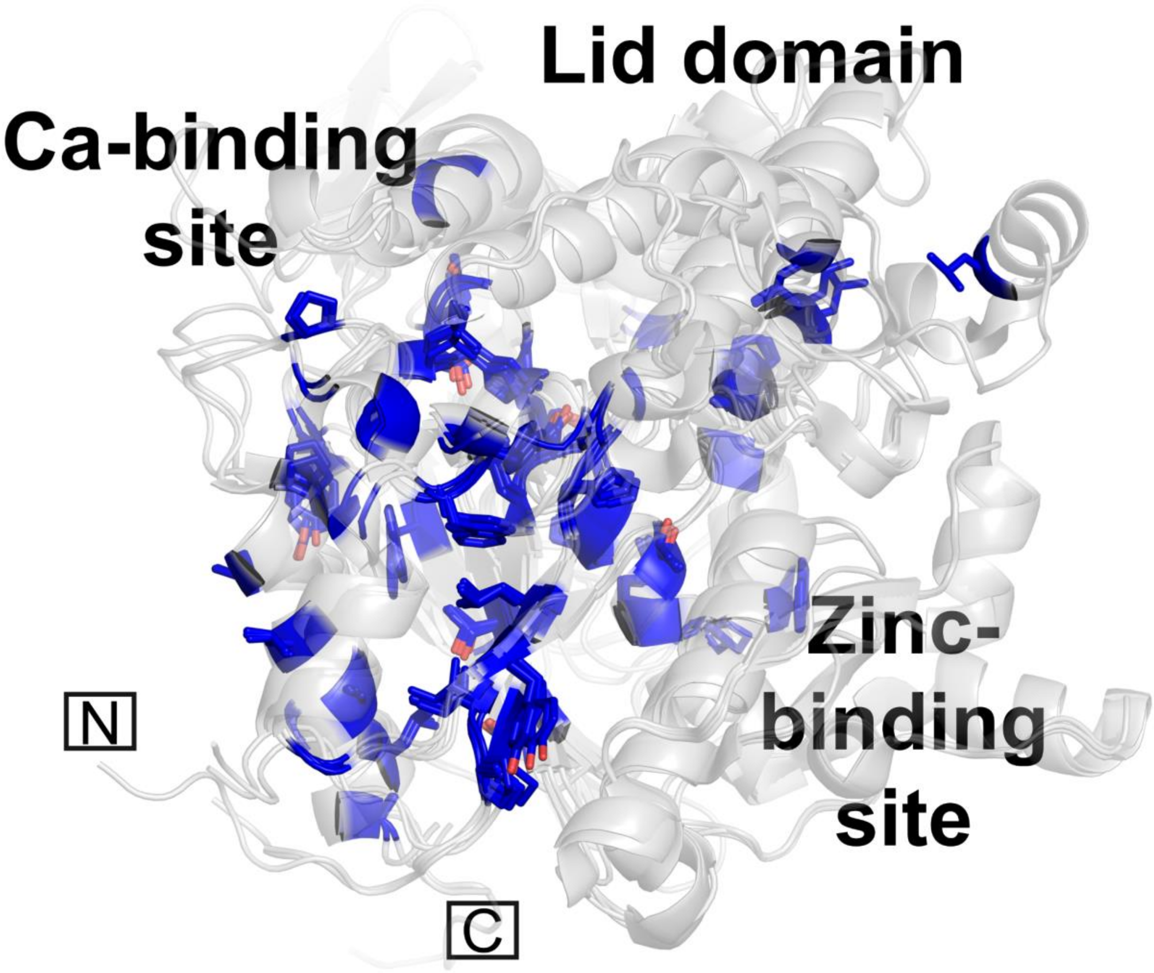

**Figure S2. Secondary-structure organization of *Gk*$_{\mathbf{Lip}}$.** Cartoon representation of the *Gk*$_{Lip\_P212121}$ structure showing the organization of the α/β-hydrolase fold. α-Helices (α1–α11) are shown in salmon and β-strands (β1–β9) in wheat and individually labeled. The lid domain (residues 169–239) is indicated at the top of the structure. The $Ca^{2+}$ - and $Zn^{2+}$ -binding regions are shown with the corresponding coordinating residues as sticks and the metal ions as spheres. The N- and C-termini are indicated for orientation.

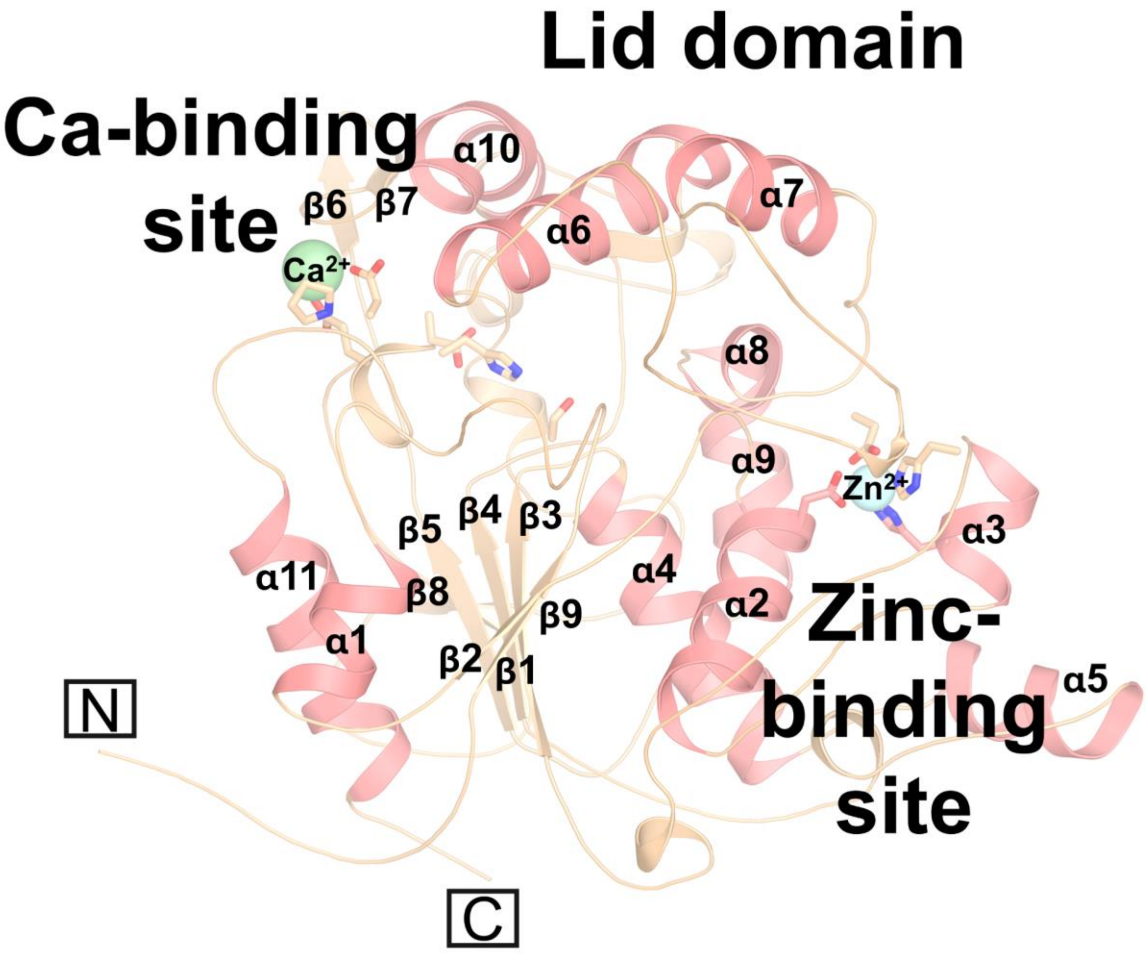

**Figure S3. Experimental electron-density support for the major functional sites of *Gk*$_{Lip}$.** *A*, overall cartoon representation of *Gk*$_{Lip\_P212121}$ highlighting the locations of the $Ca^{2+}$ -binding site, catalytic center, $Zn^{2+}$ -binding site, and lid domain, with boxed regions indicating the sites shown in detail. The N- and C-termini are indicated. *B*, close-up view of the $Ca^{2+}$ -binding site formed by Gly287, Glu361, Asp366, and Pro367. The measured $Ca^{2+}$ coordination distances are 2.3 Å to Gly287, 2.6 Å to Glu361, 2.8 Å to Asp366, and 2.4 Å to Pro367. *C*, close-up view of the catalytic triad comprising Ser114, His359, and Asp318. The indicated distances are 3.7 Å between Ser114 and His359 and 4.6 Å between His359 and Asp318. *D*, close-up view of the $Zn^{2+}$-binding site formed by Asp62, His82, His88, and Asp239. The measured $Zn^{2+}$ coordination distances are 2.7 Å to His82, 2.1 Å to His88, 2.1 and 2.7 Å to Asp62, and 2.0 and 2.7 Å to Asp239. Dashed lines indicate the corresponding interatomic distances or metal-coordination interactions.

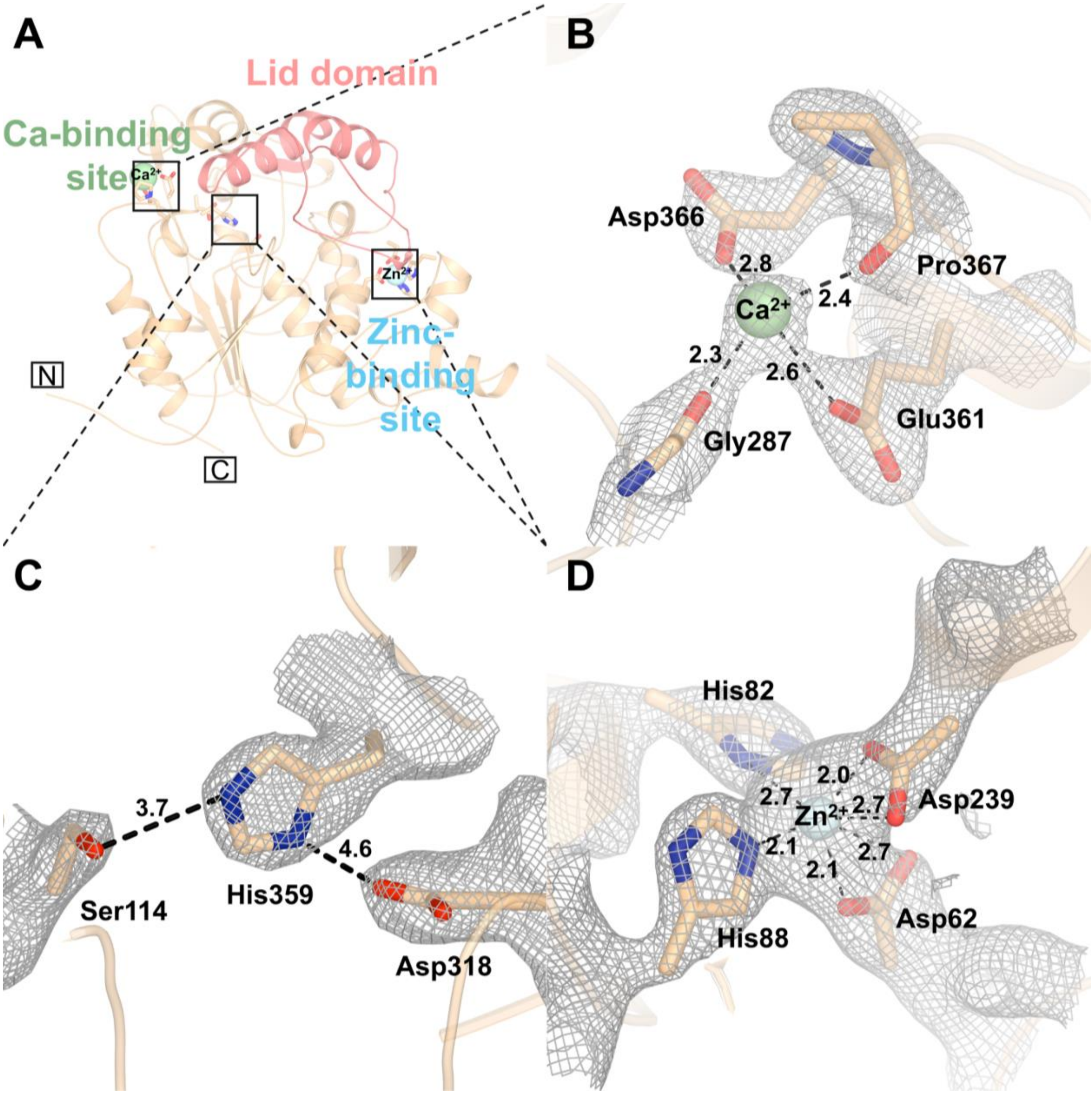


.

**Figure S4. Residue-wise structural comparison of the *Gk*$_{Lip\_P212121}$ and *Gk*$_{Lip\_P1}$ crystal forms of *Gk*$_{Lip}$ .** Per-residue backbone RMSD values between *Gk*$_{Lip\_P212121}$ and each of the four independent chains of *Gk*$_{Lip\_P1}$ (*Gk*$_{Lip\_P1A}$ –*Gk*$_{Lip\_P1D}$ ) are plotted for residues 3–389. RMSD values were calculated using matched backbone atoms (N, Cα, C, and O) following structural superposition of each *Gk*$_{Lip\_P1}$ chain onto the *Gk*$_{Lip\_P212121}$ structure. The vertical dashed lines indicate the boundaries of the lid domain (residues 169–239). The four profiles show the local structural deviations of the individual *Gk*$_{Lip\_P1}$ chains relative to the *Gk*$_{Lip\_P212121}$ reference structure.

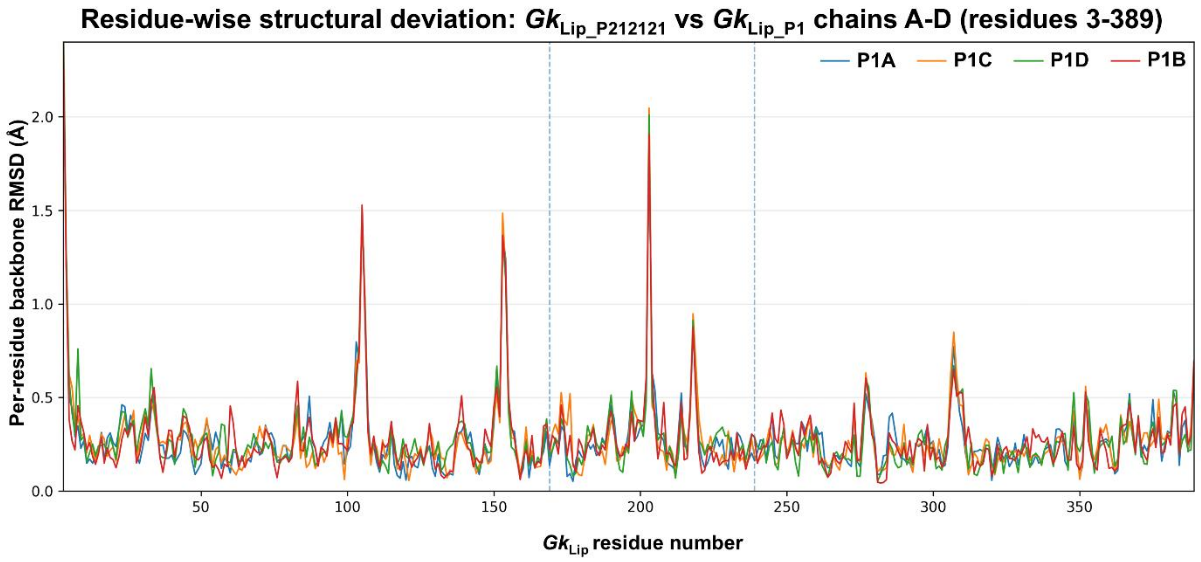

**Table 1.** ***Gk*$_{Lip}$ structure determination and refinement statistics**

| **Data Collection** | | |
|---|---|---|
| | ***Gk*$_{Lip\_P212121}$** | ***Gk*$_{Lip\_P1}$** |
| PDB ID | 45HU | 45IY |
| Beamline | P14 | P14 |
| Space group | P 21 21 21 | P1 |
| **Cell dimensions** | | |
| a, b, c (Å) | 72.86, 74.25, 102.64 | 71.50, 74.28, 102.02 |
| α, β, γ (°) | 90.00, 90.00, 90.00 | 89.97, 89.98, 90.20 |
| Resolution (Å) | 60.16–2.47 (2.51–2.47) | 102.0–2.30 (2.77–2.30) |
| CC(1/2) | 0.99 (0.48) | 0.49 (0.21) |
| I/σ(I) | 7.28 (1.49) | 2.3 (1.3) |
| Completeness (%) | 97.9 (99.5) | 75.9 (37.2) |
| Redundancy | 12.44 (12.25) | 3.6 (3.1) |
| **Refinement** | | |
| Resolution (Å) | 52.00–2.47 | 102.02–2.30 |
| No. reflections | 20,164 | 34,901 |
| Rwork/Rfree | 0.18/0.23 | 0.21/0.27 |
| No. non-hydrogen atoms | 3,300 | 12,282 |
| Protein | 3,067 | 12,232 |
| Ligand/ions | 2 | 4 |
| Water | 231 | 46 |
| **B-factors (Å$^2$)** | | |
| Protein | 37.0 | 22.1 |
| Ca2+ | 60 | – |
| $Zn^{2+}$ | 33 | 10.5 |
| Water | 39.4 | 15.9 |
| **R.m.s. deviations** | | |
| Bond lengths (Å) | 0.004 | 0.006 |
| Bond angles (°) | 0.586 | 0.87 |
| **Ramachandran plot** | | |
| Favored (%) | 97 | 97.3 |
| Allowed (%) | 3 | 2.7 |
| Disallowed (%) | 0 | 0 |

Values in parentheses correspond to the highest-resolution shell. For *Gk*$_{Lip\_P1}$, completeness values refer to ellipsoidal completeness after anisotropic cutoff and correction using STARANISO.

**Table S2. Residue-wise RMSD comparison of functional sites between the *Gk*$_{Lip\_P212121}$ and *Gk*$_{Lip\_P1}$ crystal forms of Gklip**. Residue-wise RMSD values were calculated between *Gk*$_{Lip\_P212121}$ and each of the four independent chains of *Gk*$_{Lip\_P1}$ (*Gk*$_{Lip\_P1A}$ – *Gk*$_{Lip\_P1D}$) following structural superposition. For the catalytic triad, oxyanion hole, $Zn^{2+}$ -binding, and $Ca^{2+}$ -binding residues, RMSDs were calculated using matched non-hydrogen atoms of each residue. Values are reported in Å. Mean ± SD represents the average residue-wise RMSD across the four *Gk*$_{Lip\_P1}$ chains relative to the *Gk*$_{Lip\_P212121}$ reference structure.

| Functional site | Residue | *Gk*$_{Lip\_P1A}$ | *Gk*$_{Lip\_P1B}$ | *Gk*$_{Lip\_P1C}$ | *Gk*$_{Lip\_P1D}$ | Mean ± SD (Å) |
|---|---|---|---|---|---|---|
| Catalytic triad | Ser114 | 0.309 | 0.257 | 0.149 | 0.298 | 0.253 ± 0.063 |
| Catalytic triad | Asp318 | 0.241 | 0.301 | 0.260 | 0.214 | 0.254 ± 0.032 |
| Catalytic triad | His359 | 0.314 | 0.365 | 0.308 | 0.298 | 0.321 ± 0.026 |
| Oxyanion hole | Phe17 | 0.478 | 0.274 | 0.888 | 0.460 | 0.525 ± 0.225 |
| Oxyanion hole | Gln115 | 1.461 | 0.873 | 1.964 | 1.664 | 1.490 ± 0.399 |
| $Zn^{2+}$ -binding site | Asp62 | 0.214 | 0.069 | 0.219 | 0.205 | 0.177 ± 0.062 |
| $Zn^{2+}$ -binding site | His82 | 1.180 | 1.177 | 1.180 | 1.191 | 1.182 ± 0.005 |
| $Zn^{2+}$ -binding site | His88 | 0.332 | 0.307 | 0.317 | 0.283 | 0.310 ± 0.018 |
| $Zn^{2+}$ -binding site | Asp239 | 0.380 | 0.345 | 0.564 | 0.427 | 0.429 ± 0.083 |
| $Ca^{2+}$ -binding site | Gly287 | 0.315 | 0.193 | 0.151 | 0.253 | 0.228 ± 0.062 |
| $Ca^{2+}$ -binding site | Glu361 | 0.142 | 0.136 | 0.298 | 0.128 | 0.176 ± 0.071 |
| $Ca^{2+}$ -binding site | Asp366 | 0.336 | 0.278 | 0.280 | 0.317 | 0.303 ± 0.025 |
| $Ca^{2+}$ -binding site | Pro367 | 0.521 | 0.361 | 0.717 | 0.465 | 0.516 ± 0.129 |

**Table S3. Pro367 pyrrolidine-ring geometry across the five crystallographically independent *Gk*$_{Lip}$ molecules.** The five endocyclic torsion angles (χ1–χ5) of the Pro367 pyrrolidine ring were measured for the $Ca^{2+}$-bound *Gk*$_{Lip\_P212121}$ structure and the four independent $Ca^{2+}$-free *Gk*$_{Lip\_P1}$ chains (*Gk*$_{Lip\_P1A}$ – *Gk*$_{Lip\_P1D}$). Pseudorotation phase angles and puckering amplitudes were calculated using the same atom-order convention for all structures, and each Pro367 ring was classified as Cγ-endo or Cγ-exo. The distance between Pro367 Cγ and the $Ca^{2+}$ position observed in *Gk*$_{Lip\_P212121}$ was also measured following structural superposition. For *Gk*$_{Lip\_P1A}$ – *Gk*$_{Lip\_P1D}$, these values therefore represent distances to the superposed $Ca^{2+}$ reference position, as no $Ca^{2+}$ ion is modeled in the *Gk*$_{Lip\_P1}$ chains.

| Structure | $Ca^{2+}$ occupancy | χ1 (°) | χ2 (°) | χ3 (°) | χ4 (°) | χ5 (°) | Pseudorotation phase P (°) | Pucker amplitude (°) | Assignment | Pro367 Cγ → G1 $Ca^{+2}$ reference position (Å) |
|---|---|---|---|---|---|---|---|---|---|---|
| ***Gk*$_{Lip\_P212121}$** | Present | 26.14 | -34.96 | 29.42 | -12.90 | -8.15 | 40.27 | 34.66 | Cγ-endo / DOWN | 4.32 |
| ***Gk*$_{Lip\_P1A}$** | Absent | 27.48 | -33.73 | 26.30 | -9.05 | -11.45 | 33.96 | 33.45 | Cγ-endo / DOWN | 4.86 |
| ***Gk*$_{Lip\_P1B}$** | Absent | 27.16 | -33.66 | 26.43 | -9.28 | -11.12 | 34.50 | 33.34 | Cγ-endo / DOWN | 4.59 |
| ***Gk*$_{Lip\_P1C}$** | Absent | -18.45 | 32.59 | -33.23 | 22.62 | -2.68 | 238.12 | 34.79 | Cγ-exo / UP | 5.67 |
| ***Gk*$_{Lip\_P1D}$** | Absent | 25.89 | -32.92 | 26.53 | -10.34 | -9.68 | 36.71 | 32.61 | Cγ-endo / DOWN | 4.76 |

Note: χ1–χ5 are the five endocyclic torsion angles of the Pro367 pyrrolidine ring. Pseudorotation phase and amplitude were calculated using the same atom-order convention for all structures. The $Ca^{2+}$-free *Gk*$_{Lip\_P1}$ crystal form contains three Cγ-endo conformers (*Gk*$_{Lip\_P1A}$, *Gk*$_{Lip\_P1B}$, *Gk*$_{Lip\_P1D}$) and one Cγ-exo conformer (*Gk*$_{Lip\_P1C}$). Distances for *Gk*$_{Lip\_P1A}$ – *Gk*$_{Lip\_P1D}$ were measured from Pro367 Cγ to the $Ca^{2+}$ position observed in *Gk*$_{Lip\_P212121}$ after independent structural superposition of each *Gk*$_{Lip\_P1}$ chain onto the *Gk*$_{Lip\_P212121}$ reference structure; no $Ca^{2+}$ ion is modeled in the *Gk*$_{Lip\_P1}$ chains.
